# Greater TMS-evoked frontoparietal effective connectivity is correlated with better cognitive performance

**DOI:** 10.1101/2020.08.03.234518

**Authors:** Timothy P. Morris, Maria Redondo-Camos, Gabriele Cattaneo, Didac Macia, Javier Solana-Sanchez, Goretti Espanya-Irla, Selma Delgado-Gallén, Vanessa Alviarez-Schulze, Catherine Pachón-Garcia, Emiliano Santarnecchi, Ehsan Tadayon, Recep Ozdemir, Jose Ma Tormos Muñoz, David Batres-Faz, Alvaro Pascual-Leone, Mouhsin M. Shafi

**Affiliations:** Berenson-Allen Center for Noninvasive Brain Stimulation, Department of Neurology, Beth Israel Deaconess Medical Center and Harvard Medical School, Boston, United States; Guttmann Brain Health Institute, Institut Guttmann, Institut Universitari de Neurorehabilitació Adscrit a la UAB, Badalona, Spain; Center for Cognitive and Brain Health, Northeastern University, Boston, United States; Universitat Autònoma de Barcelona, Bellaterra, Spain; Hinda and Arthur Marcus Institute for Aging Research and Center for Memory Health, Hebrew SeniorLife, Department of Neurology, Harvard Medical School, Boston, United States; Departament de Medicina, Facultat de Medicina i Ciències de la Salut i Institut de Neurociències, Universitat de Barcelona, Barcelona, Spain

**Keywords:** TMS-EEG, Frontoparietal, Cognition, Spatiotemporal dynamics

## Abstract

Fronto-parietal activity has been related to fluid intelligence and flexible cognitive control. However, causal insights on this relation are lacking. We used real-time integration of MRI-guided TMS and EEG to characterize the spatial and temporal properties of signal propagation between these two regions and relate them to cognitive performance.

31 healthy adults (55 ±6 years, 20 female) underwent TMS-EEG and a full cognitive assessment. Local and propagated current from 5 source space-reconstructed scouts ipsilateral to two stimulation sites (pre frontal cortex (PFC) and inferior parietal lobule (IPL)) was quantified in two-time windows (15-40ms and 40-80ms) and related to domain-general (global cognition) and domain-specific (memory, working memory, reasoning, flexibility, lexical access and visuo-spatial) cognitive functions.

TMS-evoked activity from stimulation of the PFC and the IPL resulted in local and distributed activity across frontoparietal regions. TMS-evoked activity in local regions was not correlated with cognitive functions. In response to TMS of the PFC, propagated current to the distal superior parietal scout in the first 15-40ms was significantly associated with global cognition (β = 2.63, SE = .898, *p* = .008, R^2^ = .31). Similarly, following TMS of the IPL, propagation to the middle prefrontal gyrus scout (15-40ms) was significantly associated with global cognition (β = 2.67, SE = 1.289, *p* = .025, R^2^ = .27). In an exploratory step, domain-specific correlations were seen in the PFC condition.

Locally evoked activity measured via source space reconstruction from TMS of two association hubs is not associated with cognitive functions. However, the propagation of the TMS pulse through frontoparietal connections is associated with overall cognitive ability. These associations are driven by a number of cognitive domains in the PFC stimulation condition.

## Introduction

The prefrontal cortex (PFC) extends projections to almost all sensory, motor, neocortical and subcortical structures and is often implicated in “top-down” modulation of cognitive functions (Miller, 2000). That is, in a hierarchical model of the neurophysiology of the cortex, the PFC constitutes the highest area of cortical representations (as opposed to lower cortical structures such as sensory and motor areas), dedicated to the integration and execution of higher-order executive functions (Fuster, 2001). Cognitive control however is not executed by a single brain region, rather via several largely non-overlapping networks, including the frontoparietal, cingulo-opercular and salience networks (Marek & Dosenbach, 2018). The frontoparietal network in particular has been implicated in numerous higher order cognitive functions such as visual working memory (Quintana & Fuster, 1999), fluid intelligence (Tschentscher et al., 2017) and task switching (Rossi et al., 2007). This relationship between frontoparietal activity and cognitive function could be because local processing of information within both the prefrontal and the inferior parietal association hubs is required for computation of higher-order cognitive functions (Culham & Kanwisher, 2001; Miller, 2000; Rossi et al., 2007). Alternatively (or in addition), the communication and interactions between these two hubs might be particularly critical for cognition; consistent with this, prior studies have also suggested that the efficiency of the connectivity and synchrony between frontal and parietal regions is associated with overall cognitive ability (Sheffield et al., 2015) and flexible cognitive control (Fries, 2015; Palva & Palva, 2011). However, the relative contribution of local processing versus inter-regional communication to cognitive function remains unclear.

Many techniques to study brain networks hinge on non-directional statistical interactions, such as in resting state functional-connectivity magnetic resonance imaging (rs-fcMRI) (Lee et al., 2013) and in resting-state magnetoelectroencephalography (MEG) and electroencephalography (EEG) (Schnitzler & Gross, 2005). Therefore, extracted insights are often reliant upon correlational statistics or algorithm-based analyses such as Bayesian techniques (Mumford & Ramsey, 2014). Meanwhile, noninvasive brain stimulation techniques such as transcranial magnetic stimulation (TMS) combined with electrophysiological imaging (EEG) can potentially reveal novel insights into the causal contribution of cortical dynamics by assessing effective connectivity. For example, TMS-EEG has demonstrated high correlation of local dynamics evoked via TMS between frontal and parietal regions, suggesting a functional overlap between evoked activity in a distributed frontoparietal network [16]. In recent work, our group has shown that TMS to adjacent nodes within parietal cortex of different resting-state networks produces network-specific propagation that correlates with cognition (Ozdemir et al., 2020). These studies illustrate the broader conceptual framework that TMS-evoked effective connectivity from perturbation of cognitively important association hubs may potentially reveal causal relations with cognitive functions (Farzan et al., 2016; Thut & Pascual-Leone, 2009).

The aims of this current study were two-fold: firstly, to quantify the TMS-evoked activity, in source space, in and between (connectivity) local and distal regions of the frontal and parietal cortices; and secondly, to gain causal insights into the differential contribution of local (frontal or parietal) activity versus distal impact from frontal to parietal and vice-versa (i.e. role of dynamics in the fronto-parietal network) on cognitive ability.

## Methods

### Subjects

31 middle-aged adults (20 female) with a mean age ± standard deviation of 55 ± 6 years (range = 43 to 64 years) were recruited as part of the Barcelona Brain Health Initiative (Cattaneo et al., 2018) and underwent a neuropsychological assessment, a TMS-EEG visit (27 completed the PFC condition, 24 for the IPL condition) and a structural T1-weighted magnetic resonance imaging (MRI). Participants were screened and consented by a medical doctor and were excluded if there were signs or evidence of any neurological or neuropsychiatric disorders. Further exclusion criteria included any contraindications for TMS (Rossini et al., 2015) or MRI. All participants gave written informed consent before participation in any study procedures, all of which conformed to the Declaration of Helsinki for research involving human subjects. The study protocol was approved by the ethics and education committee of the Institut Guttmann (Badalona, Spain).

### Neuropsychological assessment

A battery of clinical pen and paper neuropsychological assessments was performed by each participant with a licensed neuropsychologist. The battery was made up of a total of 16 validated tasks including measures of short-term memory (Rey auditory verbal learning tasks, digit-span forward, Corsi block tapping), working memory (digit-span backward, letter-number sequencing), reasoning (Weschler-adult intelligence scale version IV (WAIS_IV) matrices and block design), lexical access (phonemic and semantic verbal fluencies), flexibility (trail making test B and B-A) and processing speed/visuo-spatial abilities (Trail making test A, WAIS-IV cancellation test, digit symbol task). Education status was defined based on the number of years spent in formal higher education. Handedness was assessed by the Edinburgh handedness questionnaire (Veale, 2014) (one participant was left-handed).

### TMS protocol

TMS was applied using a Medtronic Magpro X100 stimulator through a Cool-B65 figure-of-eight coil. 120 biphasic single pulses were applied during each stimulation block at 120% of each participant’ resting motor threshold (RMT) at random intervals between 3-6 seconds, sent automatically via a custom Python script. RMT was assessed at the motor hotspot (M1) following the recommendations from the International Federation for Clinical Neurophysiology (Rossini et al., 2015) defined as the lowest stimulation intensity required to evoke motor evoked potentials (MEPs) of ≥ 50 μV in the relaxed first dorsal interosseus in five out of ten trials. MEPs were measured using surface electromyography (EMG) with electrodes placed in a belly-tendon montage on the right FDI (target muscle) and the right abductor pollicis brevis (reference muscle) with the ground on the ulnar styloid process. Electrodes were connected to a Biopac EMG100C amplifier (BIOPAC systems Inc, California, USA).

The coil was placed tangentially over the scalp roughly at around a 45-degree angle (relative to the mid-sagittal plane) resulting in a posterior-to-anterior current flow. A frameless stereotactic neuronavigation system (Brainsight, Rogue Research Inc., Montreal, QC Canada) was used in conjunction with the subject’s own T1 weighted structural MRI (obtained from a 3T Siemens magnetom prisma) to ensure accurate targeting of the stimulation sites throughout the session. Stimulation was applied to two target sites in the left hemisphere: the superior half of the posterior third of the middle frontal gyrus approximately 3cm anterior to the precentral sulcus (dorsolateral prefrontal cortex, PFC) and the inferior parietal lobule (IPL) at the superior edge of the angular gyrus roughly 1cm inferior to the intraparietal sulcus. One investigator (TM) was responsible for defining the cortical targets for all the subjects to ensure consistency. This resulted in the following average coordinates for the various stimulation site:

1. PFC stimulation site roughly below the F3 electrode with an average ± SD MNI coordinate of x = −36.3 ± 8.5, y = 37.4 ± 10.9, z = 53.3 ± 19.
2. IPL stimulation site roughly below the P3 electrode (using the international 10-20 system) with an average ± SD Montreal neurological institute (MNI) coordinate of x = −42.3 ± 10.2, y = −67.6 ± 14.4, z = 42.1 ± 23.4

Electric field modeling of the induced electric field from each stimulation condition was performed using SimNIBS v3.1 (Thielscher et al., 2015), after creating a tetrahedral headmesh for each subject using their structural T1-weighted MRI via headreco (Nielsen et al., 2018). E-field maps were normalized by the maximum E-field for each stimulation condition and for each subject, and then mapped to the fsaverage common template and averaged across subjects for the purpose of visualization (**Figure 1A**).

**Figure 1.**
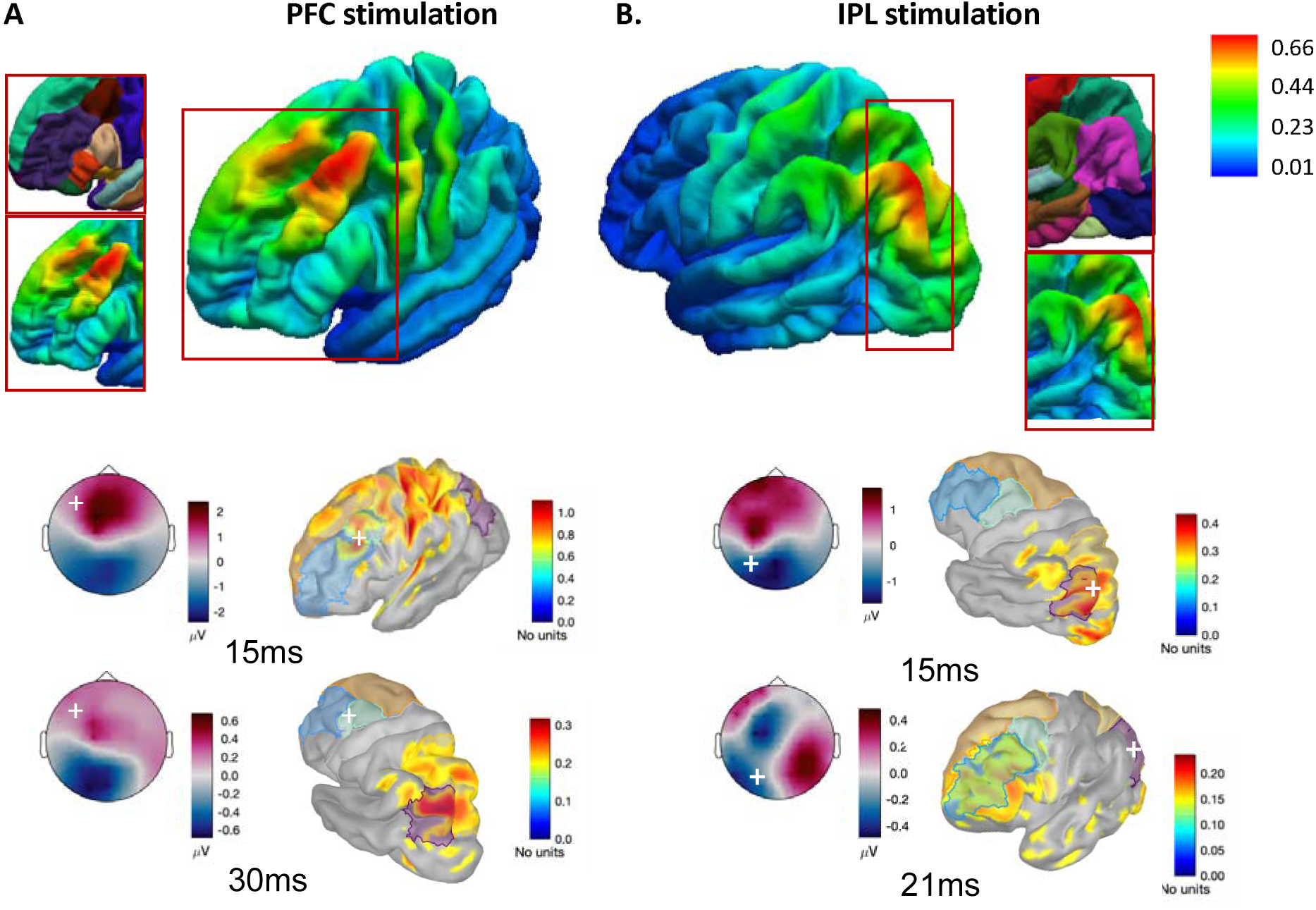
E-field modelling of the stimulation site and source space reconstructed current densities. **A and B top.** The E-field maps were normalized for each subject by the maximum E field for each stimulation condition, and then mapped to the fsaverage common template and averaged across subjects for the purpose of visualization. **A and B bottom.** EEG topoplots and source space reconstructed current density for two representative subjects showing early TMS evoked activity in the cortical region below the stimulation site (A-PFC, B-IPL) (15ms, due to the zero padding of the signal up to +14ms following the TMS pulse in the preprocessing of the EEG data). This is followed by propagation of current to the ipsilateral distal scouts in the respective parietal or frontal regions within 20-30ms after the TMS pulse. White crosses indicate stimulation site.

### EEG recordings

A TMS-compatible EEG amplifier (Brainvision ActiChamp, Brain Products, GmbH, Munich, Germany) attached to 64 active electrodes (actiCAP slim, Brain Products, GmbH, Munich, Germany) was used to record EEG responses to TMS. The ground was placed at the Fpz electrode site, and the signal was referenced to the AFz electrode. The impedance of the electrodes was kept below 5 kΩ during the recording. A continuous signal was collected, filtered at 0.1-500Hz and digitized at a sampling rate of 1000Hz. In addition to wearing earplugs to protect from the “click” of the TMS pulse, subjects listened to white noise during stimulation aimed at dampening the auditory evoked potential. The volume of the white noise was individually adjusted to each subjects’ tolerance.

### EEG preprocessing

The EEG signal was first preprocessed offline with custom MATLAB (R2017b, The MathWorks Inc, Natick, Massachusetts) scripts that incorporate the use of EEGLAB (Delorme & Makeig, 2004) and TESA (Rogasch et al., 2017). The EEG signal was epoched around the TMS pulse (−1000ms to +2000ms) followed by baseline correction with an 800ms interval (−900ms to −100ms). Excessively noisy channels were removed (with an upper limit of 2 and 3 channels deleted for IPL and PFC, respectively). Data was zero-padded between −2ms and +14ms around the TMS pulse to remove the early TMS pulse artifact. Epochs were subsequently inspected visually and excessively noisy epochs were removed (mean ± SD of # of epochs removed = 25 ± 13 for IPL and 17 ± 11 for PFC). A two-step fast independent component analysis (fICA) was performed to avoid the possibility of inaccurate ICA decompositions by large amplitude TMS-evoked muscle artefacts. Immediately prior to the first fICA, principal component analysis (PCA) was performed to reduce the data to 40 dimensions, which aimed to minimize overfitting and noisy components. Early TMS-evoked high-amplitude artefacts (either muscle or electrode) showing exponential decays were removed during the first fICA after visual inspection (with an upper deletion maximum of 3 components for both IPL and PFC). Missing data (zero-padded period of −2 to +14ms around the pulse) was linearly interpolated. A Butterworth band-pass (1Hz and 100Hz) and notch filter (48Hz and 52Hz) were applied and data was re-referenced to the average reference. A second PCA was performed to minimize the number of dimensions to 38 and after the second fICA additional artefactual components, including eye blinks, lateral eye movement, muscle, TMS-evoked muscle, electrode noise and auditory evoked potentials were removed (27 ± 3 components removed, leaving a mean ± SD of 10 ± 3 neural components per dataset for IPL and 27 ± 3 removed leaving 10 ± 4 neural components for the PFC condition). Lastly, the data was interpolated for missing channels.

### Source reconstruction

Cortical source activation was computed as current densities using Brainstorm software (Tadel et al., 2011). A three-layer boundary element model (BEM) of the head using the OpenMEEG software (Gramfort et al., 2010) was computed using individual Freesurfer-processed structural T1-weighted MRIs. Minimum norm estimates of the cortical current source density were mapped through the forward model. Because current density maps tend to preferentially extract source activity in more superficial areas of the cortex we implemented the normalization method using the dynamic statistical parametric mapping (dSPM) method (Dale et al., 2000). This normalization results in a z-score statistical map by normalizing the minimum norm estimate by the square root of the variance estimates. A constrained orientation source model was chosen (normal to cortex) which models one dipole orientated perpendicular to the cortical surface (Tadel et al., 2011). A lead field matrix was obtained by projecting standard electrode positions onto the scalp using the available ICBM152 BrainProducts ActiCap 65 electrode positions within Brainstorm. A sign flip method implemented in Brainstorm flipped the sign of the less dominant orientation before averaging (Tadel et al., 2011). For visualization purposes only, spatial smoothing using SurfStatSmooth (Surf Stat, KJ Worsely) with full width at half maximum of 3mm was performed.

Current density was extracted from *a priori* ROI’s using individual anatomical scouts of the Desikan-Killiany maps. Because we were interested in intra-frontal and intra-parietal activity as well as frontoparietal propagation and because we stimulated the left hemisphere, we chose to extract activity from five scouts ipsilateral to both stimulation conditions: the rostral middle frontal and caudal middle frontal gyri (rMf, cMf-our E-field modeling showed our stimulation site to overlap both anatomical scouts), the adjacent superior frontal gyrus (sFr), the inferior parietal lobule (IPL-the stimulation site) and its adjacent superior parietal lobule (sPa). A sixth scout, the left primary motor cortex (M1) was also used as a control condition, whereby we hypothesized that current density propagation to this scout from either the prefrontal or parietal stimulation sites would *not* be associated with global cognition. There is ongoing debate regarding the presence of non-transcranial components in TMS-evoked EEG (Conde et al., 2019) with a number of studies suggesting that later responses (>∼80ms) reflect non-specific contributions from peripherally evoked somatosensory and auditory potentials (Conde et al., 2019; Freedberg et al., 2020). Consequently, we chose to look at early evoked activity (15-80ms). The sum of the absolute values (current densities) in two time windows of interest (early; 15-40ms, and mid 41-80ms) were ultimately used as our predictor variables.

### Statistical analysis

All statistical analyses were performed in R Version 2019 (R Foundation for Statistical Computing, Vienna, Austria). Propagation to each scout of interest was quantified for each time window of interest (sum) and a one-way repeated measures analysis of variance assessed statistical differences between TMS-evoked propagation to each scout. Standardized z-scores for each of the theoretically defined (by a licensed neuropsychologist (GC)) cognitive domains were calculated and an overall cognitive ability (global cognition) composite score was created as the sum of the individual z-scores. The independent associations between source current densities and cognitive performance were assessed using separate (for each stimulation condition) multiple linear regression, controlling for both age and education level. Model assumptions were checked using Q-Q and fitted vs residual plots in R and the normality of the residuals was formally checked using Shapiro-Wilk tests of normality. The significant influence of outliers were checked using Cooke’s distance with a cut off of 0.5. Model fitness is presented as adjusted R^2^ values and significance is considered at the p = < .05 level. Multiple comparisons were corrected for using Bejamini and Hochberg’s false discovery rate (FDR) at a *q* value of 0.05, after pooling the *p* values from the regression analysis. FDR-adjusted p values (FDRpval) are presented. In an exploratory step, for those models where a significant relationship was found, we further asked whether they were being driven by any particular cognitive domain using Pearson’s correlation coefficients.

## Results

All subjects tolerated the experimental procedures – and specifically the TMS – well, and no adverse events were reported. Mean ± SD of the RMT was 56 ± 7.6 % of the maximal stimulator output intensity (MSO). Thus, mean stimulation intensity for the experiment was 68 ± 9.2 %MSO for spTMS.

### TMS-evoked propagation

No significant difference in current density across scouts in either time window of interest was found for either the PFC condition (early: F(5,156) = 0.72, *p* = .609; mid: F(5,156) = 0.783, *p* = .564) (**2A**)), or the IPL condition F(5,138) = 1.29, *p* = .268; mid: F(5,138) = 0.66, *p* = .653 (**2C**)).

The pattern of maximally-evoked activity for each stimulation condition is illustrated in two representative subjects in **Figure 1B** (and at the group level in **supplementary figure 1.** Video clips of both representative subjects and the group level are also found in supplementary material). For both participants, propagation to the respective distal frontal or parietal region is seen by ∼30ms (PFC condition (**1B** left) and ∼21ms (IPL condition (**1B** right)).

### Associations with cognitive functions

#### Local effects

Following TMS to the PFC, evoked-current density in the local scouts (cMf, rMf, sFr) were not associated (all p>0.05) with global cognitive performance in either time window of interest (see supplementary table 1 for full model results). Similarly, following TMS of the IPL, evoked-current density in the local scouts (sPa and IPL) were not associated (all p >0.05) with global cognitive performance in either time window of interest (supplementary table 1).

#### Distal (propagation) effects

In the PFC condition, propagation of current density to the distal sPa in the early time window of interest was significantly associated with global cognitive function (β = 2.64, SE = .898, *p* = .008, R^2^ = .31. **Figure 2B**), which was significant after FDR corrections (FDRpval = 0.042). This association was not seen in the other distal IPL scout (supplementary table 1).

**Figure 2.**
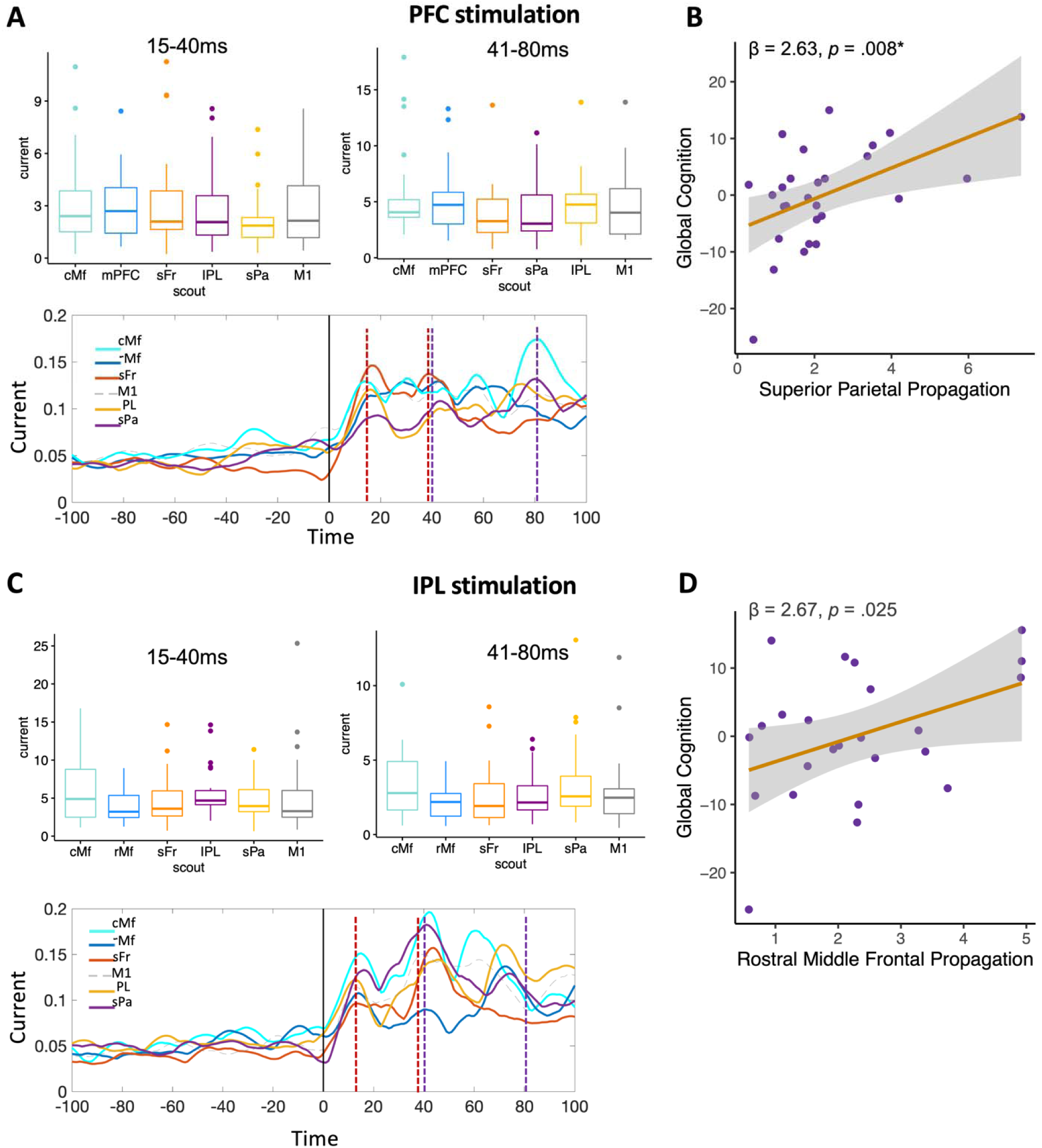
Quantification of TMS-evoked activity in source space and associations with cognition. Absolute current density in two time windows (15-40ms; 41-80ms and indicated by dashed lines time series plots) are summed for each scout. No significant differences in evoked current across scouts was found (**A- top**; **C- top**). Time series of current density in 5 source maps are plotted −100 to +100ms around the TMS pulse for the PFC (**A- bottom)** and IPL (**C- bottom**) conditions. **B and D:** Scatter plots showing the significant positive relationship between TMS-evoked propagation to distal (**B:** sPa for PFC stimulation; **C:** rMf for IPL stimulation) scouts and global cognitive ability. *indicates significance after FDR corrections.

For the IPL condition, propagation of current density to the distal mPFC scout was significantly associated with global cognitive function (β = 2.67, SE = 1.289, *p* = .025, R^2^ = .271. **Figure 2D**), in the early (15-40ms) time window of interest, however this did not survive FDR corrections (FDRpval = 0.125). Current density in the sFr and cMf scouts were not associated with global cognitive function (supplementary table 1).

#### Control condition

As a control analysis, we extracted evoked current densities from the motor cortex. TMS-evoked propagation to the motor cortex was not associated with global cognitive function in either the PFC (β = 0.956, SE = .741, *p* = .209, R^2^ = .12) or the IPL (β = 0.018, SE = .800, *p* = .983, R^2^ = .11) stimulation condition.

#### Domain-specificity

In an exploratory step, we asked whether the associations with global cognitive function from TMS-evoked propagation to distal regions were being driven by any particular cognitive domain. For PFC-evoked propagation to the sPa, short-term memory (r = 0.43, *p* = .026), lexical access (r = 0.42, *p* = .030) and visuospatial abilities (r = 0.46, *p* = .015), were all correlated with sPa current density in the early time window of interest. For IPL stimulation, no individual cognitive domain was significantly correlated with current density in the rMf in the early time window of interest.

## Discussion

Numerous studies have implicated the PFC, the parietal cortex and their connectivity (frontoparietal network) in flexible cognitive control. Nevertheless, causal inferences are difficult to establish given that most prior studies have passively evaluated brain activity without direct perturbation. With the integration of MRI-guided TMS and EEG, we characterize the spatial and temporal properties of signal propagation between the interconnected frontal and parietal regions using source space reconstruction. Here we show that early evoked propagation through frontoparietal connections significantly explains the variance in global cognitive function. In contrast, locally evoked activity was not associated with global cognition. Domain-specific effects were seen for TMS of the PFC, whereas no such domain specificity was seen with IPL stimulation.

A rich literature exists demonstrating the frontoparietal networks’ role in cognitive control, which is crucial for supporting superior cognitive functioning (Sheffield et al., 2015). The frontoparietal control network (FPCN) has been shown to have a role in control initiation, task adaptation and implementation, and providing flexible feedback control (Marek & Dosenbach, 2018). Importantly, the efficiency of this network has been positively associated with overall cognitive ability (Sheffield et al., 2015). Notwithstanding, causality in these studies is often difficult to establish given the methodology employed. Here we add to the body of literature on frontoparietal connectivity and cognition by measuring spatiotemporal cortical dynamics of effective connectivity through frontal and parietal regions using source space-reconstructed activity using MRI-guided TMS-EEG.

TMS of the frontal and parietal regions resulted in distributed activity in all of the cortical scouts of interest, suggesting that TMS stimulation of these association hubs produces activation of broadly distributed brain networks. The associations with cognition, however, were scout-specific and only seen in propagated activity rather than evoked activity in local scouts. Resting-state literature of both electrophysiological and functional magnetic resonance imaging (fMRI) have described reproducible segregated cortical networks where strong co-fluctuations of activity are seen across distributed areas of the cortex (Florin & Baillet, 2015; Thomas Yeo et al., 2011). Our results appear to show associations with cognition in the anatomical cortical regions more often associated with the FPCN (middle frontal gyrus (rMf) and superior parietal cortex (sPa) than the task-negative default mode network (superior/medial frontal gyrus (sFr), IPL) (Thomas Yeo et al., 2011).

Whilst the associations with cognitive function from IPL stimulation did not reach significance after multiple comparison corrections, the associations from PFC stimulation did. In an exploratory step we found that the association with global cognition was being driven by correlations across several cognitive tasks/domains. The fact that several domain-specific correlations are seen is consistent with the PFC role in the top-down regulation of higher-order cognitive control (Miller, 2000), as this process would be expected to effect multiple tasks.

TMS-EEG has previously shown sufficient sensitivity in the probing of fine-grained resting-state network level dynamics (Ozdemir et al., 2020), whereby the extent to which TMS-evoked activity within the dorsal attention network relative to evoked activity within the default mode network was correlated with better overall cognitive performance (Ozdemir et al., 2020). Consequently, given a number of studies have shown that the resultant physiological effect of TMS in the brain differs as a function of the stimulation site (Castrillon et al., 2020; Rosanova et al., 2009), future studies exploring the optimal methodology to target cortical networks related to cognitive function with TMS-EEG will advance our understanding of how temporal dynamics of brain function are related to cognitive performance.

There are a number of limitations to the current study. Firstly, because we stimulated the left hemisphere and were specifically interested in the relationship between cognition and frontoparietal connectivity, we focused solely on the ipsilateral hemisphere. Future analyses may focus on other networks and whole brain dynamics. Second, we did not include a sham TMS condition. Recent publications have demonstrated peripherally evoked contributions of TMS (somatosensory evoked potentials and auditory evoked potentials) that can mask the transcranial evoked cortical activity (Conde et al., 2019). However, we believe that a number of factors suggest our findings reflect transcranial evoked cortical activity. First, we employed a two-step ICA to remove TMS-specific artifacts, such as muscle, electrode, cardiac and auditory artifacts (see supplementary figure 2). Secondly, we restricted our analysis to the first 80ms. Multiple studies have empirically demonstrated site and condition specific effects of TMS-EEG at these earlier latencies, most likely reflecting transcranial evoked cortical activity (Biabani et al., 2019; Freedberg et al., 2020; Herring et al., 2015) as opposed to the non-specific peripherally evoked activity seen at later latencies (Conde et al., 2019).

TMS of the human cortex will result in propagation of activation via structural and functional connections. How these causal brain interactions relate to cognitive abilities can be explored via source space reconstruction of TMS-evoked EEG potentials. We show that rather than local cortical reactivity, TMS-evoked frontoparietal effective connectivity significantly explains domain-general cognitive ability.

## Acknowledgments

A special thanks is extended to all participants and other partners (Ad-Salutem Institute, Sodexo, I.C.A. Informa tica y Comunicaciones Avanzadas, Neuroelectrics, Corporacio Cata-lana de Mitjans Audiovisuals, Club Metropolitan, Casa Ametller, and Agència de Qualitat i Avaluacio Sanitàries de Catalunya-AQuAS) for their invaluable collaboration.

