## Supplementary material for "Greater TMS-evoked frontoparietal effective connectivity is correlated with better cognitive performance"

| S. Table 1. Associations between current density and global cognitive function | | | | | |
| --- | --- | --- | --- | --- | --- |
| **Early (15-40ms)** | β | SE | *P* | R2 | Pearson's r |
| *PFC stimulation* |  |  |  |  |  |
| cMf | 0.265 | 0.700 | 0.708 | 0.062 | 0.134 |
| rMf | 0.369 | 0.899 | 0.683 | 0.064 | 0.180 |
| sFr | 0.626 | 0.612 | 0.310 | 0.099 | 0.210 |
| IPL | 0.729 | 0.784 | 0.362 | 0.091 | 0.240 |
| ***sPa*** | 2.636 | 0.898 | 0.008* | 0.314 | 0.496 |
| M1 | 0.956 | 0.741 | 0.209 | 0.121 | 0.346 |
| *IPL stimulation* |  |  |  |  |  |
| cMf | 0.106 | 0.987 | 0.147 | 0.115 | 0.282 |
| *rMf* | 2.677 | 1.289 | 0.025 | 0.271 | 0.411 |
| sFr | -0.464 | 0.954 | 0.632 | 0.124 | -0.025 |
| IPL | 0.577 | 1.238 | 0.646 | 0.124 | 0.165 |
| sPa | 0.020 | 0.692 | 0.977 | 0.114 | -0.049 |
| M1 | 0.018 | 0.800 | 0.983 | 0.114 | 0.164 |
| **Mid (41-80ms)** |  |  |  |  |  |
| *PFC stimulation* |  |  |  |  |  |
| cMf | 0.346 | 0.467 | 0.466 | 0.078 | 0.275 |
| rMf | 0.613 | 0.603 | 0.320 | 0.097 | 0.321 |
| sFr | 0.571 | 0.652 | 0.391 | 0.087 | 0.125 |
| IPL | -0.068 | 0.684 | 0.921 | 0.057 | 0.006 |
| sPa | -0.074 | 0.579 | 0.900 | 0.057 | 0.009 |
| M1 | 0.485 | 0.570 | 0.409 | 0.084 | 0.218 |
| *IPL stimulation* |  |  |  |  |  |
| cMf | 0.176 | 0.487 | 0.720 | 0.120 | 0.195 |
| rMf | 0.374 | 0.707 | 0.603 | 0.126 | 0.140 |
| sFr | 0.208 | 0.594 | 0.730 | 0.119 | -0.072 |
| IPL | -0.379 | 0.653 | 0.568 | 0.128 | -0.017 |
| sPa | -0.367 | -0.519 | 0.609 | 0.126 | -0.029 |
| M1 | -0.163 | 0.391 | 0.680 | 0.122 | 0.037 |
| ^All models are controlling for age and education level. * Significant after FDR corrections. R2 values are adjusted R2 for the number of predictors (3); cMf = caudal middle frontal, rMf = rostral middle frontal gyrus, sFr = superior frontal lobule, IPL = inferior parietal lobule, sPa = superior parietal lobule.^ | | | | | |

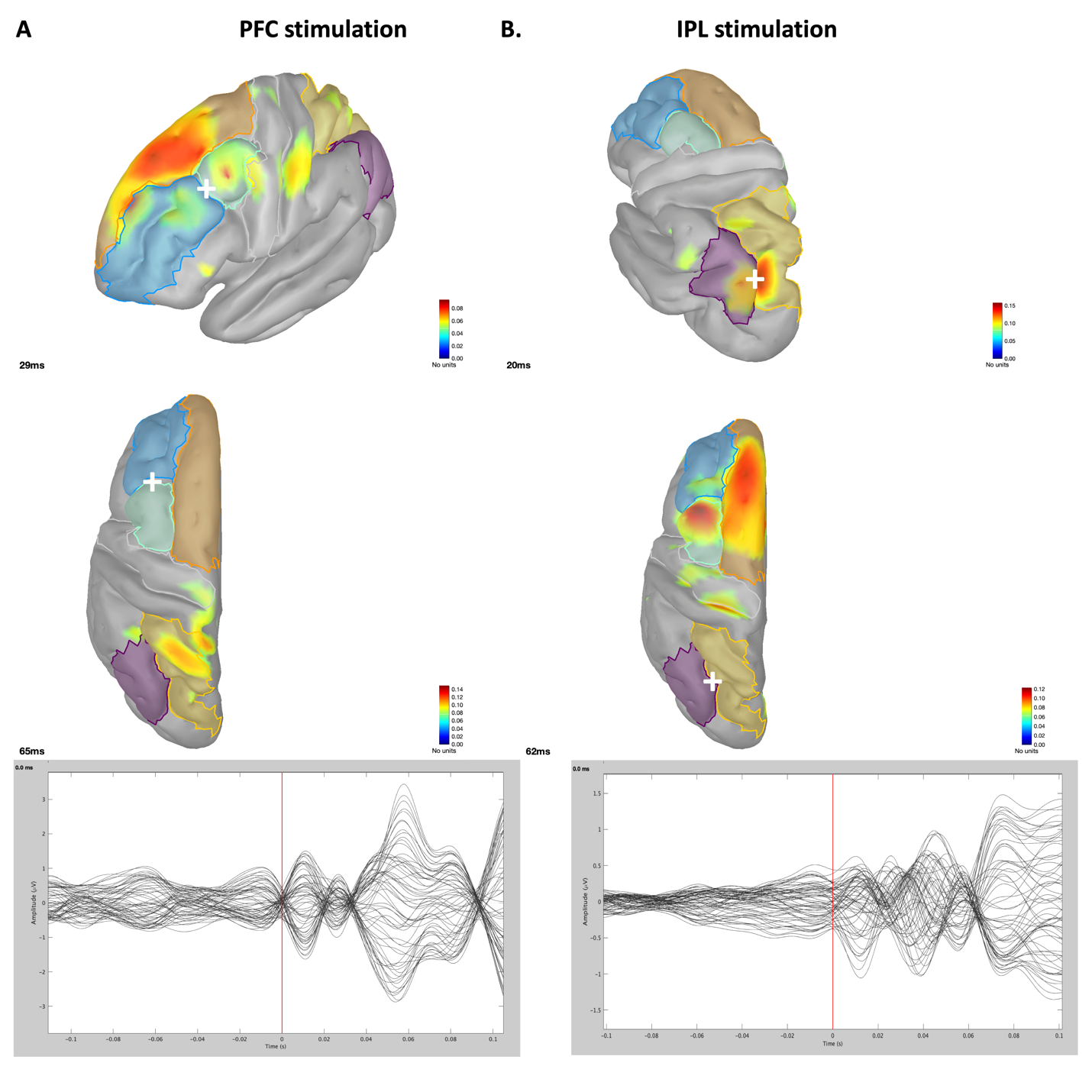
**Supplementary figure 1 caption.** Group averaged TMS-evoked activity for each stimulation condition (PFC- right; IPL- left) showing early peak activity in the stimulation site, followed by propagation to the ipsilateral distal regions by ~62ms in both conditions (at the group level). Below- group-averaged butterfly plots, -100ms to 100ms around the TMS pulse (red line) of the electrode level activity.

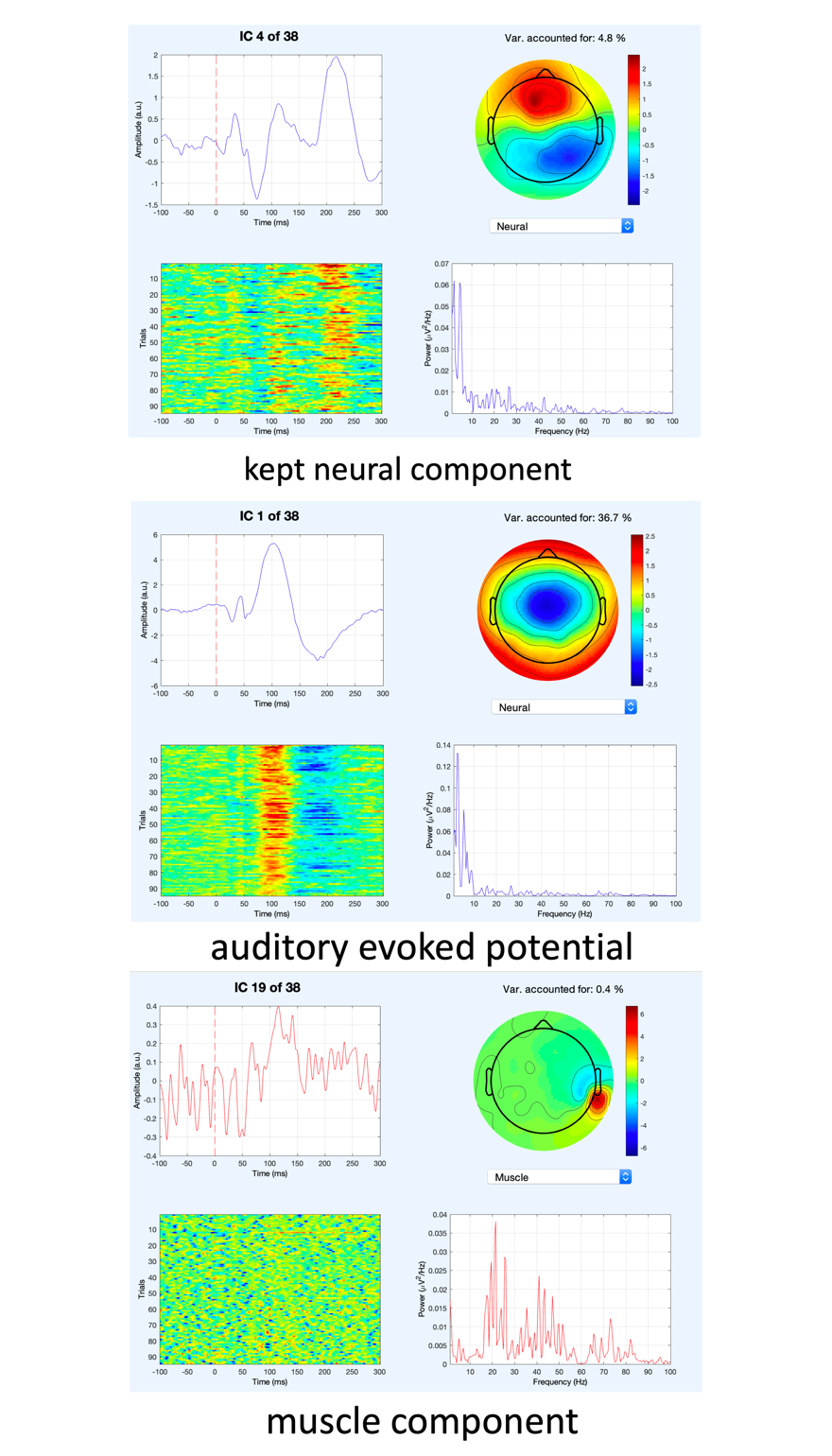

**Supplementary figure 2 caption.** A two-step ICA was applied to remove TMS-specific artifacts in the EEG signal. During the second ICA, muscle, eye blink/movements, cardiac and auditory artefacts were removed. This figure illustrates a number of examples of neural complements that were kept (top), clear auditory evoked potentials that were removed (middle) and muscle components that were removed (bottom).

**Supplementary video 1 caption.** Movie file (-20ms to +80ms around the TMS pulse) showing EEG sensor level butterfly plot (top), source space reconstructed current density with overlaid scouts of interest (bottom left) and the sensor level EEG topoplot (bottom right) for the **IPL stimulation** condition.

**Supplementary video 2 caption.** Movie file (-20ms to +80ms around the TMS pulse) showing EEG sensor level butterfly plot (top), source space reconstructed current density with overlaid scouts of interest (bottom left) and the sensor level EEG topoplot (bottom right) for the **PFC stimulation** condition.

**Supplementary video 3 caption.** Movie file (-20ms to +80ms around the TMS pulse) showing EEG sensor level butterfly plot (top), source space reconstructed current density with overlaid scouts of interest (bottom left) and the sensor level EEG topoplot (bottom right) for the **PFC stimulation condition at the group level**.

**Supplementary video 4 caption.** Movie file (-20ms to +80ms around the TMS pulse) showing EEG sensor level butterfly plot (top), source space reconstructed current density with overlaid scouts of interest (bottom left) and the sensor level EEG topoplot (bottom right) for the **IPL stimulation condition at the group level**.
